# Clueless forms dynamic, insulin-responsive bliss particles sensitive to stress

**DOI:** 10.1101/552455

**Authors:** K. M. Sheard, S. A. Thibault-Sennett, A. Sen, F. Shewmaker, R. T. Cox

## Abstract

Mitochondria perform a myriad of biochemical functions in the cell that integrate ATP production and metabolism. While mitochondria contain their own genome, mtDNA, it only encodes thirteen proteins required for oxidative phosphorylation, thus well over one thousand proteins required for all mitochondrial functions are encoded in the nucleus. One such protein is Drosophila Clueless (Clu), whose vertebrate homolog is Clustered mitochondria homolog (Cluh). Clu/Cluh is a ribonucleoprotein that binds mRNAs destined for import into mitochondria and is an essential protein that regulates cellular metabolism. Clu forms large particles in the cytoplasm, although how these particles relate to nutrition and metabolic stress is unknown. Using live-imaging, we show Clu particles are highly dynamic. Clu particles appear to be unique as they do not colocalize with many known cytoplasmic bodies. In addition, Clu particle formation is highly dependent on diet as ovaries from starved females no longer contain Clu particles and insulin is necessary and sufficient for Clu particle formation. Oxidative stress also disperses particle. Since Clu particles are only present under optimal conditions we are naming them bliss particles. These observations identify Clu particles as unique, stress-sensitive cytoplasmic ribonucleoprotein particles whose absence corresponds with altered mitochondrial function and localization.

## Introduction

Mitochondria supply the majority of ATP to tissues. Mutations in either mitochondrial DNA (mtDNA) or nuclear genes required for mitochondrial pathways can cause mitochondrial diseases that tend to affect high energy tissues (1). Metazoan mtDNA encodes thirteen proteins, 2 rRNAs and 22 tRNAs that must be supplemented by over 1,000 products encoded in the nucleus (2). How these proteins are regulated, targeted and the effect of their loss is only understood for a subset.

One such nucleus-encoded protein is Clueless (Clu). Clu encodes a large multi-domain protein that is directly involved in regulating mitochondrial function, although many details are still not well understood (3). Loss of *clu* in Drosophila is adult lethal with flies surviving only 4-7 days post-eclosion (4,5). ATP levels are normal in larvae however levels are greatly decreased in adults who accumulate mitochondrial oxidative damage (5). *clu* mutants are male and female sterile and mitochondria are clumped and mislocalized in female germ cells, which contain damaged, swollen mitochondria (3). Drosophila Clu physically and genetically interacts with the PINK1/Parkin mitophagy complex and thus may play a role as a sensor linking mitochondrial function with destruction (6). In mice, knockout of the vertebrate homolog Cluh causes post-natal lethality between P0-1, with no apparent respiratory failure, concomitant with a 10-15% decrease in weight (7). Schatton et al found post-natal death is due to metabolic disruption in hepatocytes which have mitochondrial mislocalization and are depleted of key enzymes used in catabolic energy pathways (7). Altered metabolism is also found in HeLa cells with CRISPR/Cas9 deleted Cluh, as well as mitochondrial mislocalization (8). In human development, changes in Cluh expression in preterm and term newborn umbilical cord mesenchymal stem cells corresponds with changes in the mitochondrial network and metabolic cellular metabolic changes in response to an oxygen-rich environment (9). Clu’s role in regulating mitochondrial function and metabolism is due to its role in mRNA regulation. Clu, Cluh and yeast Clu1p are ribonucleoproteins (RNPs) (4,7,10). Cluh binds and regulates mRNAs encoding proteins that will be imported into mitochondria (7,10). Clu and Clu1p bind mRNA in Drosophila and yeast, respectively, and Clu was shown in Drosophila to associate with the ribosome, potentially at the mitochondrial outer membrane (4). All of these observations underscore Clu’s fundamental role in regulating mitochondrial function, metabolism and mitochondrial protein levels through its role as a ribonucleoprotein.

While molecular analysis has clearly demonstrated Clu is an RNP, Clu also has a characteristic subcellular pattern and it is not currently clear how this relates to its function as an RNP. Clu forms large particles in Drosophila female germ cells that are closely juxtaposed with mitochondria (3). Particles are also found in larval neuroblasts and other neuronal cell types and larval muscle (5,11). Cluh forms particles, along with the Arabidopsis thaliana homolog, *friendly mitochondria* (*FMT*), which like Drosophila Clu is found in close proximity to mitochondria (8,10,12). How Clu particles relate to mitochondrial function, and how they change with metabolic or nutritional changes is not known.

Here, we examine the dynamic control of Clu particle formation and disaggregation to show Clu forms a unique and highly stress-sensitive cytoplasmic particle. Live-imaging shows Clu particles are dynamic and require an intact microtubule cytoskeleton in order to move processively. The oocyte does not contain particles and has very low levels of Clu protein relative to the connected nurse cells. Clu particles do not colocalize with other well described cytoplasmic components, thus we believe these particles are unique. Yeast Clu1p also forms particles, and *clu1* deletion causes decreased growth on a non-fermentable carbon source. In addition, Clu particles are highly sensitive to nutrition and insulin. Starved follicles no longer have particles in germ cells and surrounding somatic follicle cells, even though Clu protein levels remain the same. This effect is at least partly regulated by insulin, as insulin is both necessary and sufficient for particle formation. Nutritional stress is not the only particle disruptor, as increased mitochondrial oxidation also causes particles to disperse. Because Clu particles are unique, and unlike stress granules and processing bodies that disperse with stress, we are naming them bliss particles. Finally, we show mitochondrial localization in germ cells is completely dependent on stress-free conditions, as any of the aforementioned stressors cause clumping and mislocalization. These observations shed light on how Clu’s subcellular localization is highly dependent on the cell’s response to insulin signaling and nutrition, which could underlie Clu’s control of mRNA.

## Results

### Clu particles are abundant and highly dynamic

Clu protein is abundant and forms particles in the cytoplasm of many cell types. Female germ cells have been an excellent tissue in which to study Clu particles as they are highly metabolically active, are very large and have a large number of particles (13). To better understand Clu’s role in the cell, we used live-imaging to dissect Clu particle dynamics. To do this, we imaged Clu::GFP in the GFP trap line *clueless^CA06604^* (14). Clu particles are more abundant using live-imaging compared to fixed tissues (Movie 1). This may be due to the smaller particles being washed out during fixation, as well as particle movement making it easier to distinguish smaller particles from background. Clu shows a mix of stochastic and directed movement. At any given time, approximately 10% of the particles appear to move in a directed manner over the course of 50 seconds (Fig. 1A-A″, yellow arrows, Movie 1). For particles that move quickly, kymographic analysis supports that the movement is microtubule based (Fig. 1B, C) (15). Adding the microtubule destabilizer colcemide causes particles and mitochondria to remain stationary and disrupts the microtubule cytoskeleton as expected (Movie 2, 3, 4, Fig. S1). The dynamics of the particles (percentage in motion at any given time, speed and switching between phases of directed motion and becoming stationary) is similar to that observed for mitochondrial motion (15). The rest of the particles under normal conditions are either fairly stationary or appear to move with the bulk cytoplasmic streaming (16). Abundant Clu particles are present in the surrounding somatic follicle cells, but do not appear to move as much, likely due to the restrictive size of the cells (Fig. 1D, Movie 5). In addition, Clu protein levels are very low in the oocyte relative to the nurse cells, and we never observe Clu particles in the oocyte (Fig. 1E). This may be due to the metabolic differences between the oocyte and nurse cells. It also suggests that Clu is not moving into the oocyte from the nurse cells during follicle development, or that it is actively degraded in the oocyte during these stages. In addition, with our fixed tissue and antibody labeling, it was not completely clear how great of a contribution the “background” Clu played. With the live-images, it appears that Clu particles comprise the majority of visible protein.

**Figure 1.**
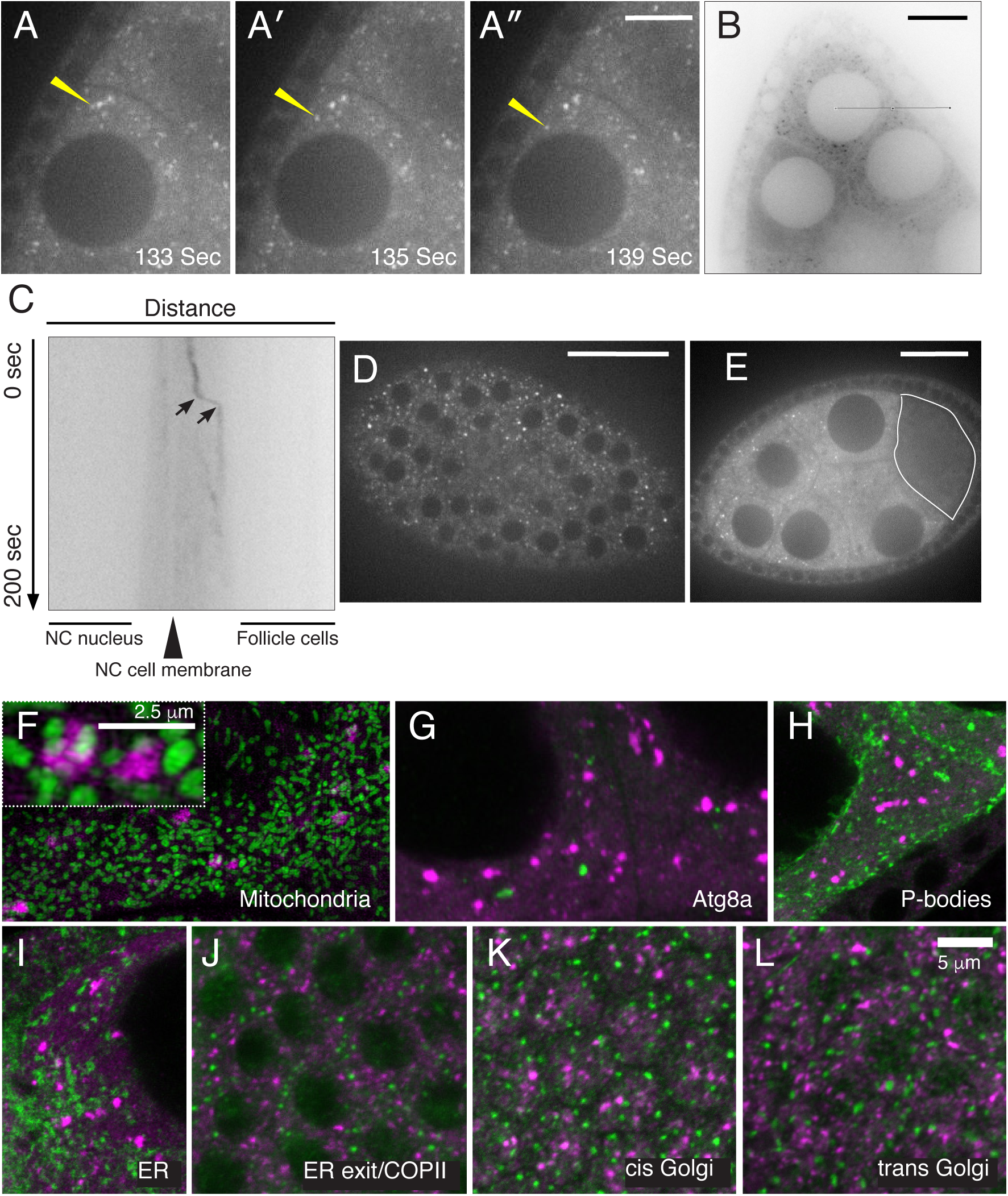
Clu forms unique, dynamic particles. (A-A′′) Follicle from *clu^CA06604^* (Clu-GFP) female. Clu particles are plentiful. Yellow arrow indicates example of processive movement. (B, C) A subset of particles move at speeds consistent with movement along microtubules. (B) Representative reverse image showing a single still-frame. The black line shows the orientation and plane used to make the kymograph (B). (C) Directed movement takes place between the arrows of the kymograph. (D) Particles are abundant in the surrounding somatic follicle cells. (E) Cross section of Clu-GFP follicle. Clu is decreased in the oocyte (white line) and does not contain particles. (F) Structured illumination micrograph. Clu particles are granular and touching mitochondria in germ cells (inset). (G, H) Clu particles do not co-localize with autolysosomes (G) or Processing bodies (H) in germ cells. (I-L) Clu particles do not appear to associate ER in germ cells (I), nor with ER-exit sites (J), cis-Golgi (K) or trans-Golgi (L) in follicle cells. Details of n values and analysis are in the materials and methods. White = Clu::GFP (A-A), Black = Clu::GFP (B), Green = ATP synthase (F), Atg8a (G), Tral::GFP (H) Sec61*α*:GFP (I), Sec23 (J), Gmap (K), and Golgin245 (L). Magenta = Clu (F-L). Scale bar: 20μm in A for A-A″, 40μm in B or B, D for D, E for E, 5 μm in L for F-L, 2.5μm in F for inset.

### Clu particles do not colocalize with many known structures

Lack of Clu causes mitochondrial deficits in all tissues examined. Since Clu interacts with several proteins on the mitochondrial outer membrane and human homolog Cluh binds mRNAs destined for mitochondrial protein import, these deficits are clearly a direct effect of protein loss (4,6,10). As we have previously shown, Clu particles tightly associate, but do not co-localize, with mitochondria (Fig. 1F)(3). Using structured illumination microscopy, the particles appear less amorphous compared to traditional confocal microscopy (Fig. 1F, inset). In fixed tissues every large particle associates with several mitochondria, but many more mitochondria are not associated. Initially, we thought this mitochondrial association may be due to Clu being involved in autophagosome formation and mitophagy. However, Clu does not co-localize with Atg8a, the LC3 homolog (Fig. 1G). Nor does Clu co-localize with a second stress-associated cytoplasmic body, processing bodies (Fig. 1H. Finally, in Drosophila larval muscle, it appeared that Clu associated with ER-exit sites (17). However, in female germ cells and surrounding follicle cells we do not observe any co-localization with components of the secretory pathway. Clu particles do not show any particular association with endoplasmic reticulum (Fig. 1I). In addition, in the follicle cells Clu particles are distinct from ER exit sites/COPII vesicles (Fig. 1J), cis Golgi (Fig. 1K) and trans Golgi (Fig. 1L). Therefore, we believe Clu forms unique particles in the cytoplasm specific to mitochondrial function.

### Clu particles are conserved

Clu has many homologs, including Cluh in humans, and we demonstrated expressing Cluh rescues the Drosophila mutant (6). The mouse Cluh homolog was shown to share many attributes of Drosophila Clu, and a Cluh knockout mouse is lethal postnatally (7,10). Yeast has an uncharacterized homolog, Clu1p (Fig. 2A). Clu1p was identified as a ribonucleoprotein by the Parker lab in a global screen for RNPs (18). The *clu1*:*CLU1*-*GFP* strain contains a GFP insertion at the endogenous *clu1* locus (19). Fixation and antibody labeling shows that Clu1p, similarly to Clu, is punctate in the cytoplasm (Fig. 2B, arrows). We have previously shown Drosophila Clu associates with the ribosome at the mitochondrial membrane and sediments in heavier fractions on a sucrose gradient, but shifts to lighter fractions with the addition of RNAse and EDTA (4). Yeast Clu1p exhibits a similar pattern on a sucrose gradient (Fig. 2C) compared to Drosophila (Fig. 5E). Since Clu directly affects mitochondrial function, we tested whether a *clu1Δ* knockout yeast strain shows defects associated with loss of mitochondrial function. Wild type yeast normally use fermentation and grow well in a fermentable carbon source such as glucose (Fig. 2D). *clu1Δ* grows equally well as wild type on media containing glucose (Fig. 2D). However, when grown on the non-fermentable carbon source glycerol, which forces the cells to use oxidative phosphorylation, *clu1Δ* shows decreased growth compared to wild type (Fig. 2D). This was true in two different wild type backgrounds (BY4741 and W303) and with three re-derived *clu1*::*KANMX* deletions. The poor growth and small colonies can be seen after growing a small number of cells on glycerol for over a week (Fig. 2E). Petite colony formation also reveals defects in genes involved in the respiratory chain and occurs in mutants that are defective in oxidative phosphorylation even when grown on a fermentable carbon source. A strong mutation will give many petite colonies. Newly derived *clu1Δ* in a BY4741 background had a slightly higher percent (6.5%) petite formation vs wild type BY4741 (3.9%) when grown on YPdextrose.

**Figure 2.**
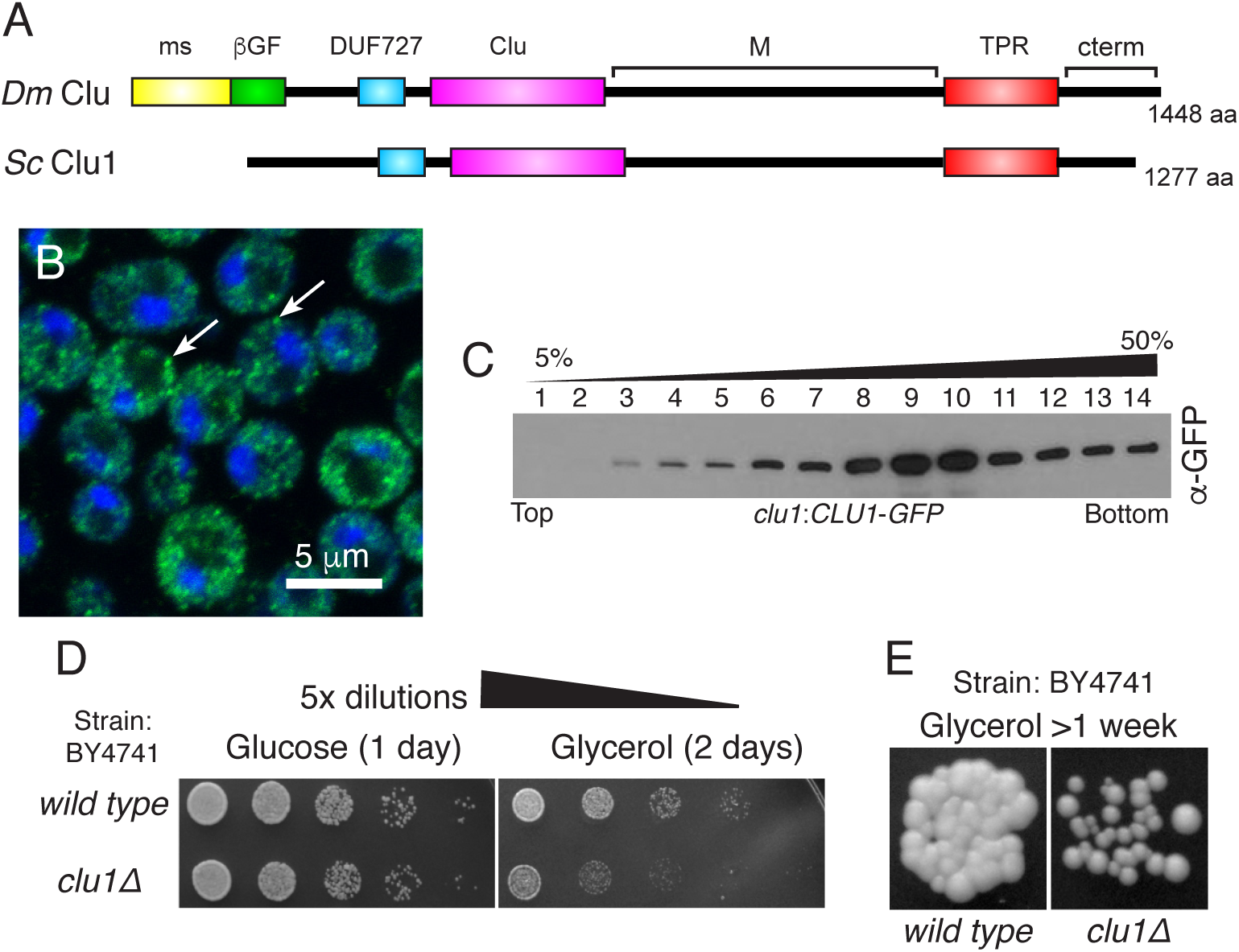
Yeast Clu1p accumulates as cytoplasmic particles. (A) Cartoon diagram of the shared domain structure between Drosophila Clu (Dm) and yeast Clu1p (Sc). (B) Fixed log-phase yeast cells of strain *clu1::Clu1-GFP*. Clu1p puncta indicated by arrows. (C) Sucrose gradient using extract from strain *clu1*::*CLU1-GFP*. Clu1p sediments in heavy high sucrose fractions. (D) Serially diluted *clu1Δ* cultures grow normally on glucose, but not on glycerol. (E) *clu1Δ* forms small colonies after one week of growth on glycerol compared to its parental wild type strain. Green = Clu1p-GFP, blue = DAPI. Scale bar: 5 μm in A.

### Clu particles are sensitive to nutritional stress

We have observed only well-fed females reproducibly have robust Clu particles, suggesting ample access to food is important. To analyze the effect of nutrition on Clu particle dynamics, we starved and refed flies in order to determine how feeding affected particle formation. Wild type females fed wet yeast paste changed daily for 4-7 days always have robust Clu particles (Fig. 3A-A″). However, starving the same well-fattened flies for five hours on water completely abolishes particles, resulting in what appears to be slightly elevated diffuse Clu in the cytoplasm (Fig. 3B-B″). Re-feeding yeast paste for as little as two hours completely reverses this effect (Fig 3C-C″, G). These results were the same for the surrounding somatic follicle cells (Fig. 3D-F″). The mild five-hour starvation (typical starvation lasts >24 hours (20,21)) does not result in any behavioral defects and levels of ATP remain the same, with a small increase in the starved flies presumably due to compensatory metabolic mechanisms (Figs. 4H, 5C). Particle disaggregation is not due to protein degradation as Clu protein levels remain the same from ovary extract of all three conditions (Fig. 3I, J). These results support nutrition as an important regulator of Clu particle formation, and that their formation and disaggregation are highly dynamic and reversible.

**Figure 3.**
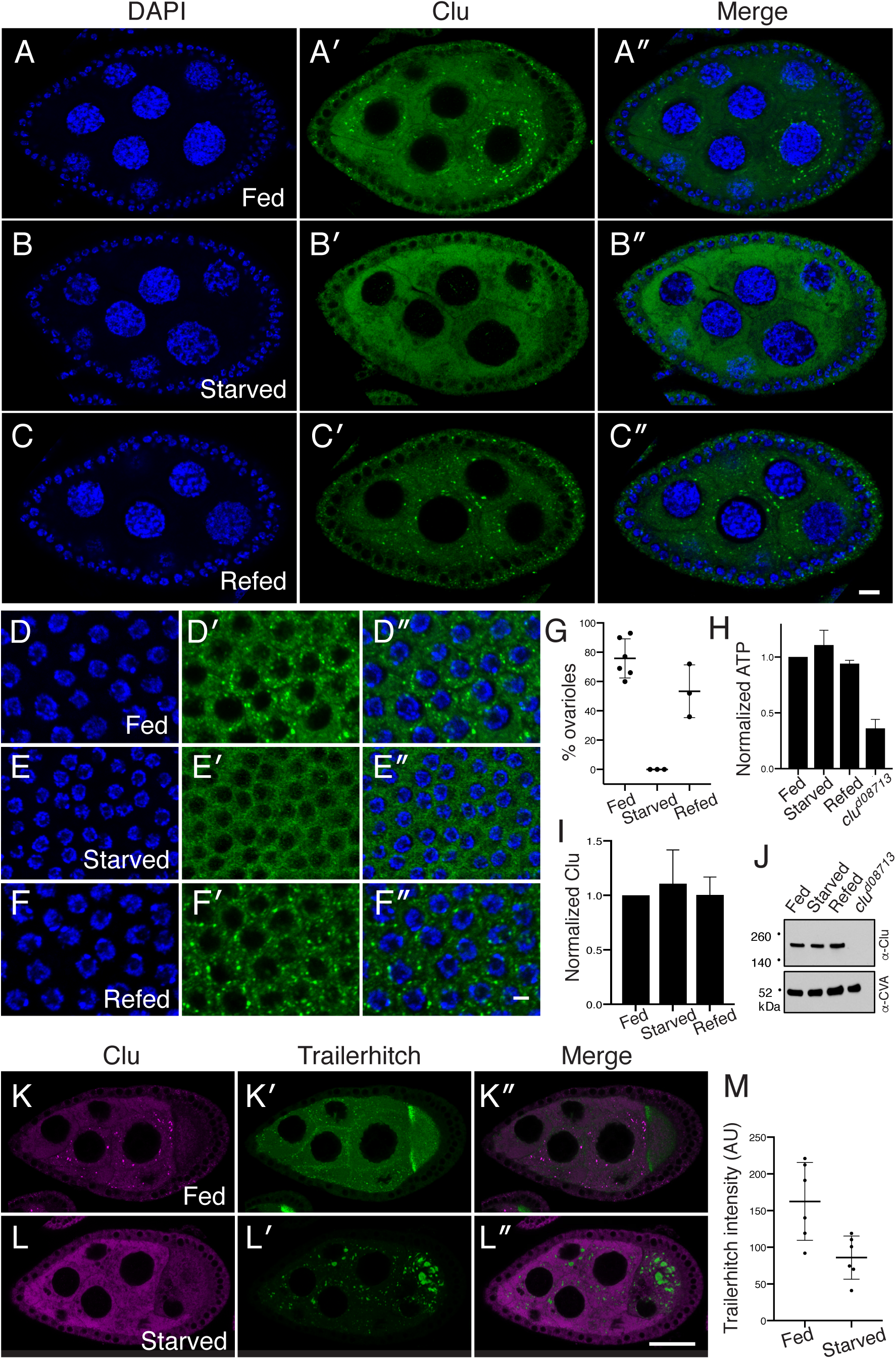
Clu particles disaggregate in response to starvation. (A-C″). Follicles from *w^1118^* females. Well-fed follicles contain many large particles in the germ cells (A). Starvation for 5 hours on H_2_0 causes the particles to disaggregate (B′). Re-feeding starved females causes Clu particles to reform (C′). Quantification is shown in G. (D-F″) Surrounding follicle cells show the same dynamic as the germ cells. ATP levels (H) and Clu levels (I, J) remain the same. *clu^d08713^* is a positive control. (K-L″) Tral::GFP has small aggregates in the nurse cells with a homogeneous background and concentration at the anterior of the oocyte (K′). Upon starvation, Tral::GFP forms very large Processing bodies and the diffuse Tral signal decreases (L′, M). Clu particles disaggregate and Clu becomes diffuse (L). Details of n values are in the materials and methods. Error bars are S.E.M. Green = Clu (A-C″), Tral::GFP (K-L″), magenta = Clu (K-L″), blue = DAPI (A-F″). Scale bar: 10 μm in C″ for A-C″ and in F″ for D-F″, 40μm in L″ for K-L″.

**Figure 4.**
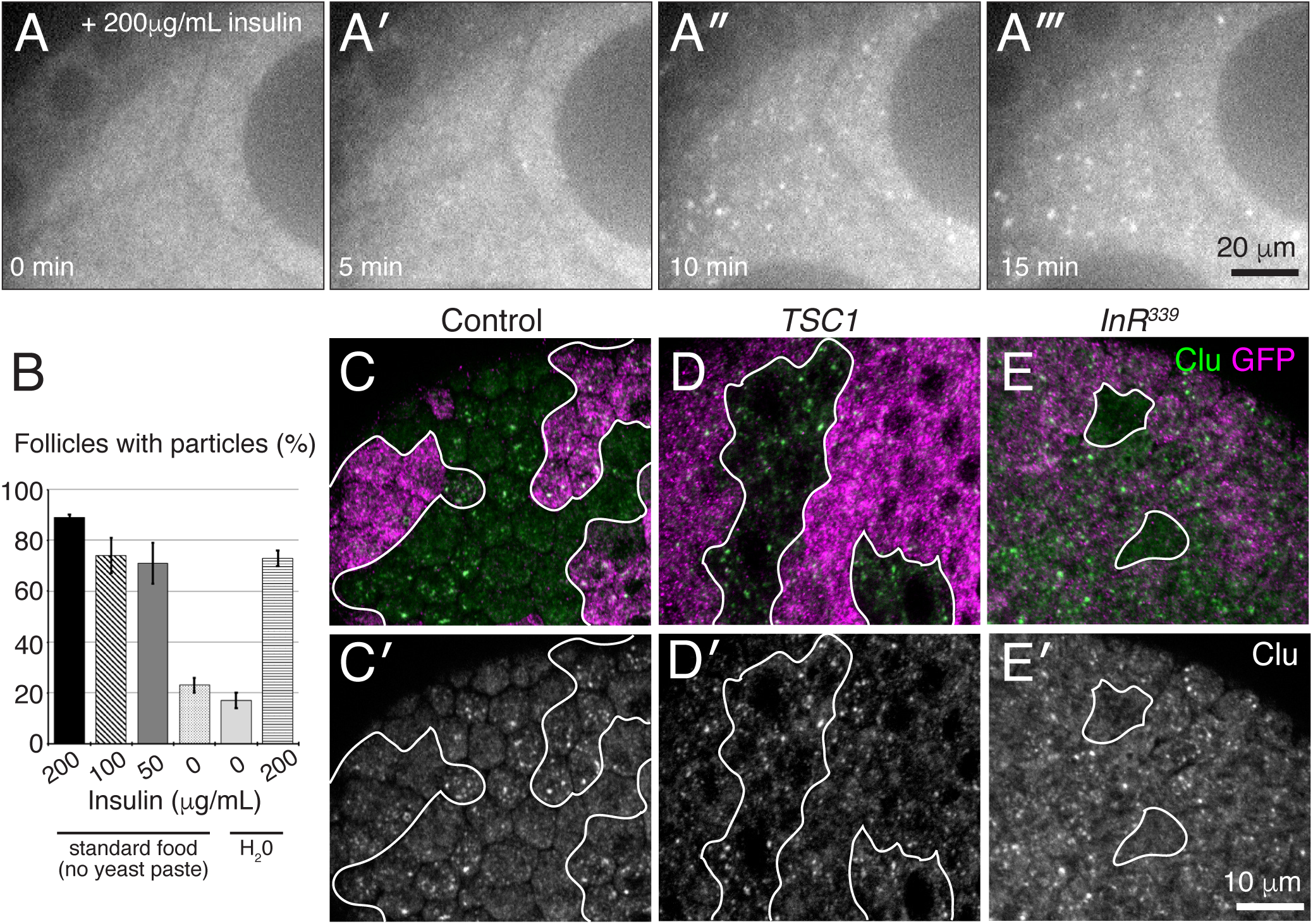
Insulin is necessary and sufficient for Clu particle formation. (A-A‴) Stills from live-image of *clu^CA06604^* germ cells. Females were raised on standard media (no yeast paste) and dissected in Schneider’s. With insulin addition, particles start forming in five minutes. (B) Quantification. (C, E′) Clonal analysis in follicle cells. Wild type control clones have Clu particles (GFP-) (C, C′). Mutant clones for *TSC1* also have Clu particles (D, D′). Mutant clones for *InR* lack particles (white circles) (E, E′). White = Clu::GFP (A-A‴), green = GFP-mutant cells, magenta = GFP positive wild type siblings. n values described in materials and methods. Error bars = SD. Scale bars: 20 μm in A‴ for A-A‴, 10 μm in E′ for C-E′.

**Figure 5.**
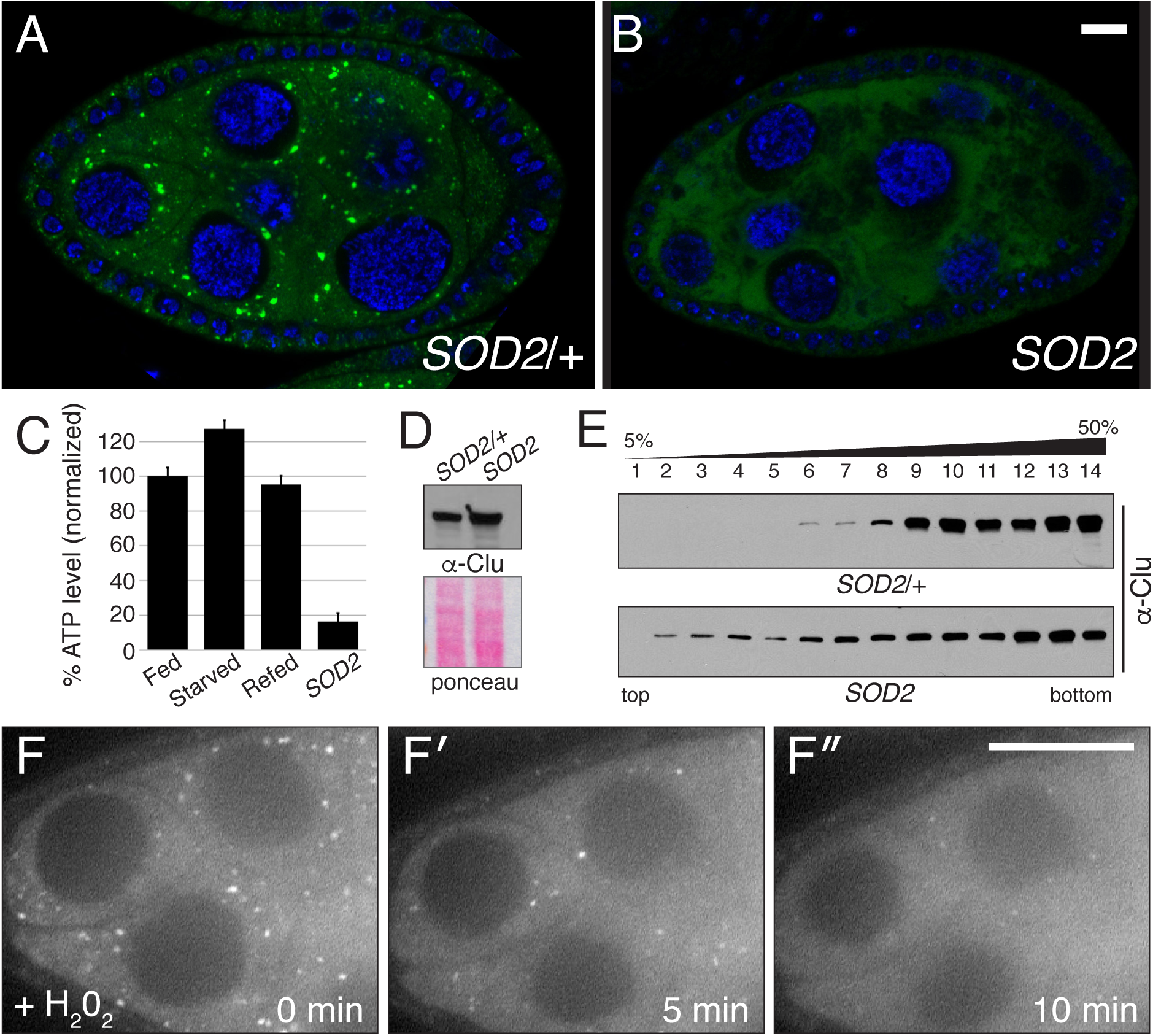
Oxidative stress disperses Clu particles. (A, B) Clu particles are missing in *SOD2* mutant follicles (B) vs wild type siblings (A). (C, D) *SOD2* mutants have decreased levels of ATP (C), but not decreased Clu protein (D). (E) A sucrose gradient of adult fly extract. *SOD2* mutant flies show a shift to lighter fractions. (F-F″) Live-image stills of a well-fed *clu^CA06604^* follicle. Addition of H_2_0_2_ causes particles to disperse. Green = Clu, blue = dapi (A, B), white = Clu::GFP (F-F″). Error bars are S.E.M. Scale bar: 10μm in B for A, B, 40μm in F″ for F-F″.

Trailerhitch (Tral) is a ribonucleoprotein that is a component of processing bodies (22,23). Under well-fed conditions, Tral::GFP forms characteristic small particles in the cytoplasm, is concentrated in the anterior of the oocyte and is also diffusely cytoplasmic (Fig. 3K′) (23). In the same follicle, Clu forms large particles (Fig. 3K). After five-hour starvation, Tral::GFP forms very large processing bodies in the nurse cells and oocyte which capture all the diffuse cytoplasmic staining causing a reduction in “background” intensity (Fig. 3L′, L″, M) (24). In contrast, in the same follicle Clu particles disperse completely with an apparent rise in diffuse cytoplasmic label (Fig. 3L, L″). These observations support that Clu particles are uniquely different from particles that form in response to nutritional stress, such as stress granules and processing bodies.

### Insulin is necessary and sufficient to induce Clu particle formation

Given how feeding affects Clu particle formation, we examined the insulin signaling pathway to determine whether this pathway was responsible for our observations. To do this, we utilized live-imaging to manipulate insulin concentration in the media. Media to image follicle maturation live was developed by the Montell laboratory (25). This media has additional insulin. Imaging follicles from *clu^CA06604^* females reared on standard food alone and dissected in media without insulin results in approximately 20% of the follicles containing Clu particles (Fig. 4B). This is similar to the number found after five-hour starvation using this assay (Fig. 4B), and both are slightly higher than dissecting in Grace’s using our standard protocol, followed by fixation (Fig. 3G). Adding insulin while imaging resulted in robust Clu particle formation within ten minutes, giving % Clu-particle-containing follicles on par with fixation (Fig. 4A-A‴, Movie 6). This result was similar whether the females were starved on water or standard food (Fig. 4B). These results support that insulin is sufficient to induce particle formation.

To further assess the role of insulin in Clu particle regulation, we used clonal analysis to increase and decrease signaling in the surrounding somatic follicle cells. In control clones containing no mutations and distinguishable by GFP (FRT82B), particles are present in both GFP+ and GFP-sibling clones (Fig. 4C, C’). To increase insulin signaling, we made clones mutant for *TSC1*, a negative regulator of insulin signaling. *TSC1* GFP-mutant clones contained particles (Fig. 4D, D’). The particles may be larger and more plentiful with this increased signaling, but this was difficult to quantitate. To determine the effect of lack of insulin signaling, we made mutant follicle cell clones of *InR* using two alleles, *InR^399^* and *InR^E19^*. Loss of *InR* resulted in a loss of Clu particles in the mutant clones compared to wild type sibling clones (Fig. 4E, E’). These results support that increasing insulin signaling has no effect on Clu particle formation, and loss of insulin signaling causes loss of particles, indicating insulin is required for particle formation.

### Clu particle formation is sensitive to mitochondrial oxidative stress

Nutritional stress and lack of insulin causes particles to disaggregate. To determine whether oxidative stress has the same effect, we examined *Superoxide Dismutase 2* (*SOD2*) mutants. SOD2 is a free-radical scavenger in the mitochondrial matrix (26). Loss of SOD2 causes an increase in mitochondrial oxidation (5,27). *SOD2* mutant flies eclose at normal numbers and appear relatively healthy at first, but die within 24 hours (5,27). Before that, their ovaries appear to develop relatively normally for this short duration, however, they completely lack Clu particles (Fig. 5B). As we have shown previously, *SOD2* mutant adults have low levels of ATP (Fig. 7C) (5), but Clu levels are not reduced (Fig. 5D). We have previously shown Clu sediments in the heaviest fractions on a sucrose gradient, and these fractions can be shifter lighter with the addition of RNAse or EDTA (4). Examining sedimentation of Clu in *SOD2* mutants vs. *SOD2*/+ control, we see there is a shift to the lighter fractions as well (Fig. 5E). This could be related to lack of Clu particles, or to a change in Clu’s role as a ribonucleoprotein or its role in binding the ribosome. To determine the effect of oxidation on particle stability using live-imaging, we added H_2_0_2_ to the insulin-containing culture media of follicles dissected from *clu^CA06604^* well-fed females. Particles started to disaggregate in as quickly as five minutes after addition of H_2_0_2_ indicating general oxidative stress is also effective (Fig. 5F-F″, Movie 7).

### Mitochondrial mislocalization in Drosophila female germ cells is downstream of stress

We have shown mitochondria mislocalize in female germ cells due to a variety of mitochondrial dysfunction, including loss of *clu*, *PINK1* and *parkin* (3,6). Under well-fed conditions, mitochondria are evenly dispersed in developing follicles in the ovary, as previously described (Fig. S2A) (28). However, after five-hour starvation, mitochondria clump in the nurse cell cytoplasm, reminiscent of *clu* mutants (Fig. S2B, arrow, F). After two hours re-feeding, mitochondria evenly disperse and are back to the well-fed pattern (Fig. S2C). *SOD2* mutants show a similar mitochondrial clumping (Fig. S2E, arrow). To use live-imaging, we added TMRE, a cell-permeant cationic dye that preferentially sequesters in mitochondria with high membrane potential. At time zero, TMRE-labeled mitochondria are evenly distributed (Fig. S2G, Movie 8). Once the follicles incubate in H_2_0_2_ for five minutes, mitochondria start to clump in the cytoplasm (Fig. S2G′). After ten minutes, oxidative damage is to the point where most mitochondria are quite dim, with only a small subset still fluorescing (Fig. S2G″). These results indicate that once the cells undergo stress, whether from oxidative damage, nutritional stress, or mutation, mitochondria no longer retain their normal location. This effect occurs quite quickly, and with respect to nutrition, can be readily reversed.

## Discussion

Here we show that Clu bliss particles are dynamic cytoplasmic bodies whose formation and dispersal is highly dependent on stress. Clu bliss particles do not co-localize with many other cellular components supporting that they are a new cytoplasmic particle specifically supporting mitochondrial function (Fig. 2). In addition, there is no localization with Atg8a, a marker for autophagosomes, which engulf damaged mitochondria during mitophagy (29) and conditions that would be expected to increase mitophagy, such as loss of *SOD2*, do not increase the number of particles (Fig. 7). Furthermore, Clu particles have the opposite response to starvation stress as processing bodies (24). The dynamic movement of particles in the germ cells is similar to other RNP particles, with particles moving at a speed consistent with mitochondrial movement (15).

Clu particles are acutely and reversibly sensitive to nutrition. Flies reared with yeast paste have large particles, which using live-imaging are more abundant than we previous realized since fixation eliminates or masks the smaller ones. A five-hour starvation completely eliminates all particles and a brief re-feeding restores them. Furthermore, insulin addition to cultured follicles causes particles to form quickly. Standard food alone or water starvation does not dramatically adversely affect the flies as they behave normally and have normal levels of ATP. Thus, the metabolic effect of lack of yeast paste quickly regulates Clu particles presumably by affecting the metabolic state of germ cells.

Drosophila Clu and vertebrate Cluh are ribonucleoproteins and Clu particles could be sites of translation for Clu-bound mRNAs. Since there are many small Clu particles present, it is possible that a larger fraction of mitochondria is associated with particles than we can see using fixed tissue imaging. Clu associates with several mitochondrial outer membrane proteins, including TOM20 and Porin, and also self-associates, however it is unclear how Clu particles associate with mitochondria (6). Attempts to simultaneously visualize mitochondria and Clu particles live have been stymied by follicle sensitivity to TMRE addition (the Clu particles disperse), though there is evidence that Clu particles in Arabidopsis do move with mitochondria (12). Clu particle formation and dispersal may be important for Clu’s role in controlling metabolism through regulating catabolic enzymes. Since the particles respond so quickly to nutritional cues, they may represent an additional control for mRNA translation, with translation rates different in the presence and absence of particles. This hypothesis remains to be tested.

There have not yet been patients presenting with mitochondrial disorders identified who harbor mutations in *Cluh*. This is somewhat surprising given how large the protein is, however, it may be that any perturbations in Cluh are either not compatible with life or cause very early lethality. This is supported by the mouse Cluh knockout and the observation of the difference in umbilical cord mesenchymal stem cells between preterm and term births in humans (7,9). Nonetheless, given how Clu forms dynamic cytoplasmic particles highly sensitive to stress and is an important RNP critical for directly affecting mitochondrial function and biogenesis, understanding Clu’s molecular role and how its mRNA binding is related to cytoplasmic localization will be important in the future.

## Materials and Methods

### Fly stocks

The following stocks were used for experiments: *w^1118^*, *clueless^d08713^*/CyO Act GFP (3), *trailerhitch^CA06517^*, *Sec61α^CC00735^*, *clueless^CA06604^* (14), *SOD2^Δ2^*/CyO Act GFP, FRT82B, FRT82B GFP/TM3 (Bloomington Drosophila Stock Center), FRT82B *TSC1^Q87X^*/TM3 (30), FRT82B *InR^339^*/TM3, FRT82B *InR^E19^* (31). Flies were reared on either standard cornmeal fly media or standard cornmeal fly media supplemented with yeast paste at 22° or 25° C.

### Clone generation

0-24hr females of the appropriate genotype were fed yeast paste for 24 hr, then heat shocked 2x 1 hour in a 37℃ water bath, morning and evening. Except for during heat shock, the flies were kept undisturbed on a quiet shelf at room temperature. After 24 hours feeding yeast paste, 3x ten flies were dissected in parallel with controls. Clones were distinguished by absence of GFP antibody labeling. *TSC1* mutant clones were abundant as the cells divide at a normal frequency. *InR* clones were very small and found at a very low frequency due to the shortened heat shock regimen. We found 10 clones for *InR^339^* and 10 clones for *InR^E19^*. Standard procedures for clone induction (3x heat shock, 7-10 days on yeast paste before dissection) gave *InR* germline clones, but did not show consistent, reproducible Clu particles for controls. Thus, the results were not interpretable, and we did not use this protocol. We assume Clu particle loss in the controls using this protocol was due to stress.

### Fixed Image Immunofluorescence

Flies were collected up to 1 day after eclosion and fattened with yeast paste for 3-7 days. 60 female flies were then transferred to an empty vial containing a Kimwipe soaked in water at the bottom of the vial for five hours to induce starvation. 30 female flies continued feeding on yeast paste as controls. 30 starved flies were then transferred back to a vial with fresh fly food and yeast paste for two hours for re-feeding. 3x 10 flies were then dissected at room temperature (RT) in Grace’s Insect Medium (modified) (BioWhittaker, Lonza, Cologne, Germany). Ovaries were fixed for 20 minutes in 4% paraformaldehyde and 20mM formic acid solution (Sigma) made in Grace’s. Tissues were washed two times (40 minutes each) with antibody wash buffer (1X PBS:0.1% Triton X-100:1% BSA) and were incubated in primary antibody over night at 4°C. They were then washed two times (40 minutes each) and incubated overnight at 4°C in secondary antibody. After washing, DAPI (1:1000) was added for five minutes then removed, then Vectashield (Vector Laboratories, Inc.) was added. For yeast staining, log phase Clu1p::GFP yeast grown in YPD were treated 20% paraformaldehyde to a final concentration of 4% for 20 min. The cells were spun, washed and spun twice with 1.2M sorbitol, 50mm KPOH, when resuspended in 500 uL of the same buffer. The cells were treated with zymolase for 35min at 37℃, then very gently washed 2x with Ab wash, followed by 2 hours *α*-GFP sitting at room temperature with occasional gentle inversion, washed 2x with Ab wash, then added and *α-*rabbit Alexa-488 for 1 hour. 2x gentle wash with dapi addition in the second wash followed. The cells were then mounted in Vectashield and imaged using a Leica AF6000 Time-lapse Imaging System. The following primary antibodies were used: guinea pig anti-Clu N-terminus (1:2000) (3), mouse anti-Complex V (ATP synthase (CVA)) (1:1000, Mitosciences, Inc, cat# MS507), rabbit anti-GFP (1:2000, AbCam, Catalog #Ab290), GABARAP (Atg8a) (1:200, AbCam, cat #Ab109364), Sec23 (1:500), Gmap (1:1000, DSHB), Golgin245 (1:1000, DSHB). The following secondary antibodies were used: goat anti-mouse IgG_2b_ Alexa 568 (1:500) and goat anti-guinea pig Alexa 488 (1:1000) (Molecular Probes, Invitrogen), donkey anti-guinea pig Cy3 (1:1000) and donkey anti-rabbit Alexa 488 (1:500) (Jackson Labs). Samples were imaged using a Zeiss 700 confocal microscope and 63x Plan Apo NA 1.4 lens.

### Image Quantification

For Clu particle quantification, slides were scanned at 40x using a Zeiss Axio Scan.Z1 slide scanner. The number of ovarioles per slide was counted (90-100 per slide) using the Cell Counter tool in ImageJ. Each ovariole on a slide was subsequently scored using the Zen blue software; an ovariole was scored as having particles if at least one follicle in the ovariole contained particles and as having no particles if no follicles contained particles. The mean percentage of ovarioles with particles in each experiment (n = 6 groups of 10 flies for well-fed group, n = 3 groups of 10 flies for starved group, n = 3 groups of 10 flies for refed group) is represented in the graph with error bars representing mean ± SD. Fluorescent intensity for Tral::GFP was calculated using equal regions of interest between starved and fed and ImageJ to calculate the mean fluorescent intensity. Kymographs were generated using ImageJ from single still-frames from videos of *clueless^CA06604^* well-fed follicles taken at 60x, with one frame captured every second for 200 seconds. Thirty-nine particles were analyzed from 12 follicles. A representative image was used in the figure.

### Live Imaging Microscopy for Drosophila Tissue Samples

*clueless^CA06604^* females (express GFP-tagged *Clu*) that had been either fattened with yeast paste or fed only the standard cornmeal fly media, were dissected at room temperature (RT) in either Complete Schneider’s Media (Schneider’s *Drosophila* media [Thermofisher] with 15% Fetal Bovine Serum and 0.6X Pen-Strep solution) or Complete Schneider’s Media supplemented with 200 µg/mL of insulin (insulin which did not readily dissolve in media was allowed to settle out overnight at 4°C prior to use). Ovaries were removed and further teased apart into single, isolated ovarioles. Ovarioles were then loaded onto a glass bottom 35mm dish (MatTek Corporation) and live videos were collected using a Nikon spinning disk/TIRF/3D-STORM microscope with photo-manipulation with X60 magnification at room temperature. During experiments when insulin was added to ovarioles during filming, ovarioles would be prepared as above in Complete Schneider’s Media and Complete Schneider’s Media containing 400 µg/mL insulin would be added to the dish with a 3mL syringe to dilute the sample 1:1, resulting in an insulin concentration of 200 µg/mL with 3 biological repeats. For Insulin response graph, flies were dissected in C.S. containing the amount of insulin indicated and viewed on Nikon spinning disk microscope as described above, individual follicles were counted to either have particles (if at least one nurse cell in follicle has particles) or no particles after focusing throughout entire follicle. Mean percentage of follicles with particles in each experiment is represented in graph with SE (calculated with standard deviation/square-root(3)). The number of follicles counted varied between 16 and 62 per biological repeat with the following follicles counted per experiment:

Non-fattened flies in C.S. with 200ug/mL insulin – 16, 39 and 62;
Non-fattened flies in 100ug/mL insulin – 27, 39, and 39;
Non-fattened flies in 50ug/mL insulin – 29, 27, and 26;
Non-fattened flies in 0ug/mL insulin – 34 and 16;
Fattened flies in 200ug/mL insulin – 29 and 20;
Starved flies in 0ug/mL insulin – 19 and 23;
Starved flies in 200ug/mL insulin – 29 and 26.

Colcemide treatments were performed as above with the follicles incubated in 1:100 Colcemide in Complete Schneider’s Media with 200 µg/mL insulin for 3 hours at room temperature before imaging. TMRE was used at a concentration of 1:10,000 and treated for 20 minutes, follicles were directly imaged with no washes. H_2_O_2_ experiments were performed at a concentration of 1µM and treatment lasted for 15 minutes while filming.

### Yeast Growth Assays

For percent petite colony formation, approximately 500 cells were plated on YPD. Normal (rho+) and petite (rho-) colonies were counted in two square regions of interest. For BY4741 wild type background: 483 rho+ out of 495 yeast cells (97.6%) and for BY4741 *clu1Δ* (newly made): 506 rho+ out of 584 cells (86.6%). For the glycerol growth assay, log-phase BY4741 wild type and *clu1Δ* strains were grown overnight in YPGlucose, then diluted the next morning and grown until achieving log-phase. The cells were then serially diluted on both YPglucose agar and YPglycerol agar and grown overnight. To image individual colonies, BY4741 wild type and *clu1Δ* strains were grown overnight in YPGlucose, then diluted an grown the next morning until log-phase. Approximately the same number of cells was spotted on YPglycerol and allowed to grow for a week before imaging.

### ATP Assays

Flies were collected up to 1 day after eclosion and fattened with yeast paste for 3-7 days. 30 female flies were then transferred to an empty vial containing a Kimwipe soaked in water at the bottom of the vial for five hours to induce starvation. 15 female flies continued feeding on yeast paste as control. 15 starved flies were then transferred back to a vial with fresh fly food and yeast paste for two hours for re-feeding. Flies were then homogenized in extraction buffer (100 mM Tris-Cl, pH 8.0, 4 mM EDTA, pH 8.0; 6 M guanidine hydrochloride) (3 groups of 5 flies/50 μL extraction buffer), boiled for 4 minutes, then centrifuged at 8000 g for 5 minutes at RT. The protein concentration of the samples was determined using a Bradford assay. The ATP concentration was determined using an ATP Determination Kit (Molecular Probes, Invitrogen) according to the manufacturer’s directions. 100 ml assays were performed in a 96 well plate and the luminescence was measured using a Biotek Synergy H1 luminometer. Each sample was read in triplicate. The amount of ATP was normalized against protein concentration.

### Western blotting

Flies were collected up to 1 day after eclosion and fattened with yeast paste for 3-7 days. 30 female flies were then transferred to an empty vial containing a Kimwipe soaked in water at the bottom of the vial for five hours to induce starvation. 15 female flies continued feeding on yeast paste as control. 15 starved flies were then transferred back to a vial with fresh fly food and yeast paste for two hours for re-feeding. Flies were then dissected at room temperature (RT) in Grace’s Insect Medium (modified) (BioWhittaker, Lonza, Cologne, Germany). Ovaries were homogenized in 150 uL of sample buffer and proteins were separated on 4-20% polyacrylamide gels using a standard SDS-PAGE protocol for Western blotting. After electrophoresis, proteins were transferred onto a Hybond-ECL nitrocellulose membrane (GE Healthsciences, Inc.) then soaked with Ponceau S for 10 min and rinsed as a further loading control. Sucrose gradients were performed as described (ref). Blots were exposed to the following antibodies: anti-Clu (1:20,000) (3) and anti-α-tubulin (1:5000, Developmental Studies Hybridoma Bank, University of Iowa, Iowa City). Quantification of Western blots was performed with ImageJ, and the amount of Clu protein was normalized against *α*-tubulin (n = 3).

### Sucrose gradients

Sucrose gradients for fly extract were performed as previously described (4). The modifications for yeast were the following: yeast from 5 ml YPD overnight culture was pelleted and transferred to an eppendorf tube. An equal volume of silica beads was added to the pellet plus 50ul of 1X mDXB. The tube was vortexed frequently for up to 5 minutes. 300ul 1X mDXB was added to the mixture and spun down at 2000g, twice for 5 minutes to collect the supernatant. Half the supernatant was loaded on the sucrose column.

## Acknowledgements

We would like to thank the USUHS Biomedical Instrumentation Core for imaging support and The Developmental Studies Hybridoma Bank, created by the NICHD of the NIH and maintained at The University of Iowa, Department of Biology. This work was supported by the National Institutes of Health (1R21NS085730 to R.T.C. and 1R35GM119790-01 to F.S.) and Uniformed Services University (USUHS BIO-71-3019) to R.T.C.

---

**Movie 1.** Clu live-imaging shows robust, dynamic particles in germ cells. Follicle from *clu^CA06604^* (Clu-GFP) female. Imaged one frame per second for 200 seconds. Video is 36 frames per second.

**Movie 2.** Mitochondria movement is normal without colcemide treatment. Follicle from *clu^CA06604^* (Clu-GFP) female treated with TMRE. Imaged one frame per second for 120 seconds. Video is 10 frames per second.

**Movie 3.** Mitochondrial movement ceases with treatment of the microtubule destabilizer colcemide. Follicle from *clu^CA06604^* (Clu-GFP) female treated with TMRE. Imaged one frame per second for 120 seconds. Video is 10 frames per second.

**Movie 4.** Clu particle movement ceases with treatment of the microtubule destabilizer colcemide. Follicle from *clu^CA06604^* (Clu-GFP) female. Imaged one frame per second for 120 seconds. Video is 24 frames per second.

**Movie 5.** Clu particles in surrounding follicle cells are more stationary than in germ cells. Follicle from *clu^CA06604^* (Clu-GFP) female. Imaged one frame per second for 120 seconds. Video is 24 frames per second.

**Movie 6.** Clu particle formation is induced with insulin. Follicle from *clu^CA06604^* (Clu-GFP) female. Imaged one frame per minute for 45 minutes. Video is 10 frames per second.

**Movie 7.** Oxidative stress disperses Clu particles. Follicle from *clu^CA06604^* (Clu-GFP) female. Imaged one frame per 15 seconds for 15 minutes. Video is 10 frames per second.

**Movie 8.** Oxidative stress causes mitochondrial mislocalization in Drosophila germ cells. Follicle from *clu^CA06604^* (Clu-GFP) female. Imaged one frame per 15 seconds for 20 minutes. Video is 10 frames per second.

**Figure S1.**
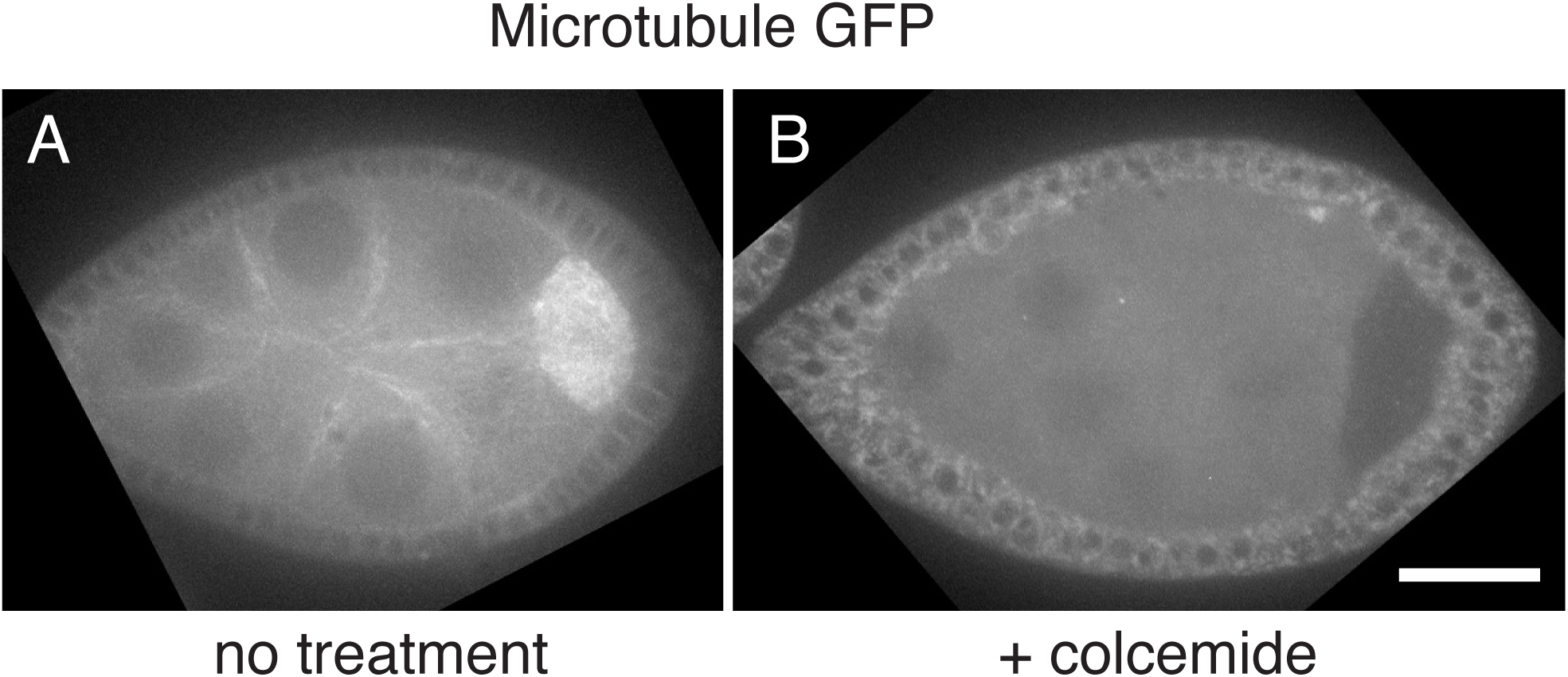
Colcemide treatment disrupts microtubules. Microtubule-GFP expressing follicles. Culturing in media shows the microtubule cytoskeleton is intact (A). Treatment with colcemide shows no microtubules (B). Scale bar: 40μm in B for A, B.

**Figure S2.**
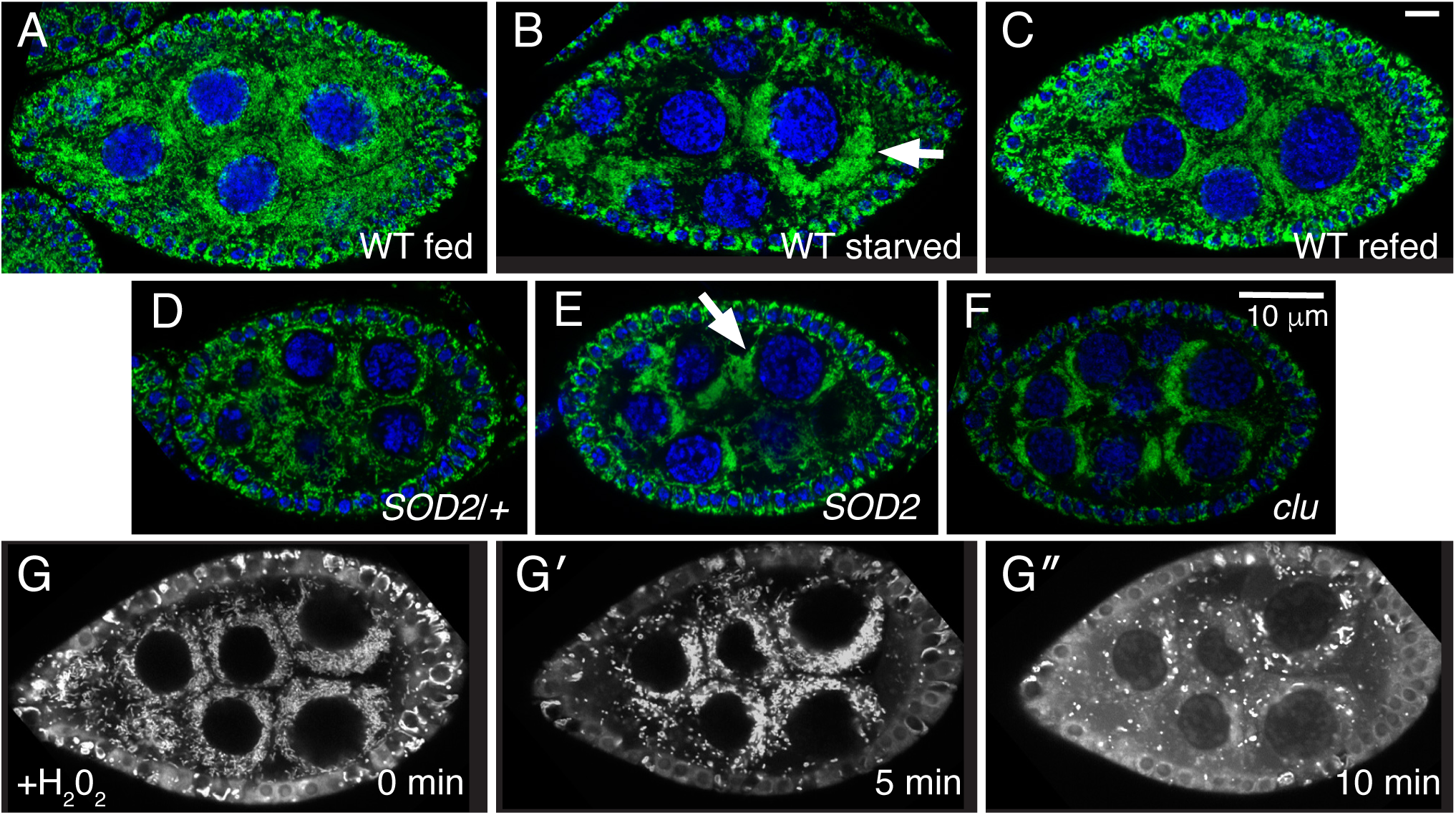
Stress causes mitochondrial mislocalization in Drosophila germ cells. (A-C) Wild type flies. Well-fed females have evenly dispersed mitochondria in germ cells (A). Starvation causes mitochondria to clump (arrow) (B). Refeeding yeast paste for two hours post-starvation causes mitochondria to disperse (C). (D, E) Mitochondria in *SOD2* mutant germ cells also form clumps (arrow) compared to wild type *SOD2*/+ siblings (D). This clumping is reminiscent of loss of Clu, as previously published (F). (G-G″) Stills from live-imaging *clu^CA06604^*. Adding H_2_0_2_ to cause oxidative stress also causes mitochondria to clump. TMRE labeling of mitochondria indicates initially mitochondria are dispersed (G), but soon after start to clump (G′). At a later time-point, the TMRE labeling becomes spotty due to mitochondria losing their membrane potential and ability to concentrate the dye (G″). Green = ATP synthase (A-F), blue = dapi (A-F), white = TMRE (G-G″). Scale bars: 10 μm in C for A-C, 10 μm in F for D-F.

